# Genome assemblies of three closely related leaf beetle species (*Galerucella* spp)

**DOI:** 10.1101/2021.04.28.441848

**Authors:** Xuyue Yang, Tanja Slotte, Peter A. Hambäck

**Affiliations:** Department of Ecology, Environment and Plant Sciences, Stockholm University, Stockholm 10691, Sweden

**Keywords:** Galerucella calmariensis, Galerucella pusilla, Galerucella tenella, Coleoptera, leaf beetle

## Abstract

*Galerucella* (Coleoptera: Chrysomelidae) is a leaf beetle genus that has been extensively used for ecological and evolutionary studies. It has also been used as biological control agent against invading purple loosestrife in North America, with large effects on biodiversity. Here we report genome assembly and annotation of three closely related *Galerucella* species: *G. calmariensis*, *G. pusilla* and *G. tenella*. The three assemblies have a genome size ranging from 460Mb to 588Mb, with N50 from 31,588kb to 79.674kb, containing 29,202 to 40,929 scaffolds. Using an *ab initio* evidence-driven approach, 30,302 to 33,794 protein-coding genes were identified and functionally annotated. These draft genomes will contribute to the understanding of host-parasitoid interactions, evolutionary comparisons of leaf beetle species and future population genomics studies.

## Introduction

*Galerucella* (Coleoptera: Chrysomelidae) is a leaf beetle genus that is distributed worldwide except in the Neotropics (Thomas *et al.* 2002). Some species have been used as biological control agents against invasive plants and the host specificity and environmental impact of these species have attracted broad interest. The most common application is the introduction of *G. calmariensis* and *G. pusilla* from Europe to North America against the invasive wetland plant purple loosestrife (*Lythrum salicaria*). Since 1992, releases of *Galerucella* populations have been made in many states in the USA and the colonization appears to have been successful, leading to a dramatic decrease of *L. salicaria* populations (Blossey *et al.* 1994; Landis *et al.* 2003; McAvoy *et al.* 2016).

In addition to its application in biological control, *Galerucella* spp. has been widely investigated in both ecological (Pappers *et al.* 2002; Tanaka and Nakasuji 2002; Hori *et al.* 2006; Fors *et al.* 2016) and evolutionary (Ikonen *et al.* 2003; Stenberg and Axelsson 2008; Yang *et al.* 2020) studies. In particular, *Galerucella* spp. has been used to study ecological and evolutionary consequences of host-parasitoid interactions (Stenberg *et al.* 2007), mainly involving three closely related species (*G. calmariensis*, *G. pusilla* and *G. tenella*) with similar life cycles and their shared wasp parasitoid (*Asecodes parviclava*) (Hambäck *et al.* 2013; Fors *et al.* 2016). The divergence of these three species is fairly recent: *G. pusilla* and *G. calmariensis* diverged around 77,000 years ago while *G. tenella* diverged around 400,000 years ago (Hambäck *et al.* 2013). *G. pusilla* and *G. calmariensis* share an exclusive host plant (*L. salicaria*), whereas *G. tenella* feeds primarily on *Filipendula ulmaria* and occasionally on other Rosaceae species. In all three species. adults in the study area overwinter until mid-May and then lay eggs on leaves or stems of their host plant. Larvae hatch after 1-2 weeks, pupate in late June to early July, and the adults emerge from the pupae by the end of July. The three species are attacked by the same endoparasitoid wasp *Asecodes parviclava*, which lays one or more eggs in the beetle larvae. The successful wasp larvae kills the host, use it as food resources, and subsequently emerge during the following summer (Hansson and Hambäck 2013).

The demography, host searching behavior and immunology have previously been addressed in several *Galerucella* species (Zheng *et al.* 2008; Fors *et al.* 2015; Yang *et al.* 2020), but no genome assemblies of *Galerucella* species are currently available. The closest related species that has an available genome assembly is ragweed leaf beetle (*Ophraella communa*), which also belongs to the leaf beetle (Chrysomelidae) family (Figure 1) (Bouchemousse et al. 2020). Here we report *de-novo* genome assemblies for *G. calmariensis*, *G. pusilla* and *G. tenella*. We performed computational annotation, assigned gene ontology to functional proteins and performed ortholog cluster analysis between the three species. These draft genomes will be useful for understanding the mechanisms underlying beetle interactions with parasitoid and plant use, and for future population genomics studies.

**Figure 1.**
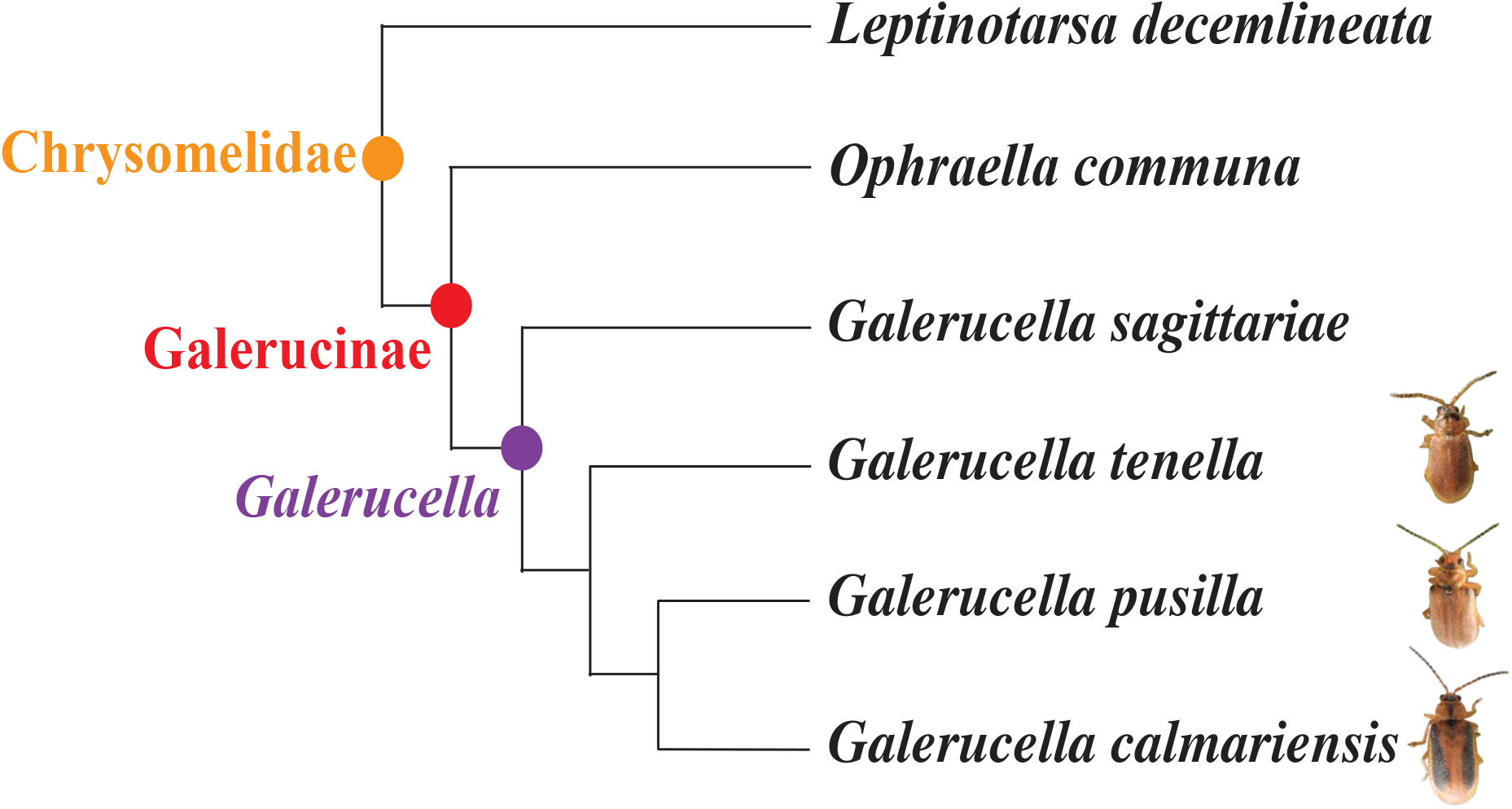
Topology of *G. calmariensis*, *G. pusilla and G. tenella* and their position in Chrysomelidae, adapted from previous studies generated by mitochondrial and nuclear genetic markers (Hambäck *et al.* 2013; Bouchemousse *et al.* 2020). The dots at nodes corresponds to the taxonomy at each branch. Length of branches is not proportional to genetic distance between clades.

## Materials and methods

### DNA extraction and sequencing

Larvae samples of *G. pusilla* and *G. calmariensis* were collected in mid-May 2018, from Iggön (59° 2′30.81“N, 17° 9′49.35“O), Sweden. *G. tenella* samples were collected in mid-May 2018, from Södersjön (59°51′8.72“N, 18° 6′26.59”O), Sweden. To reduce heterozygosity and bacterial contamination, we reared and inbred the beetles in the laboratory at room temperature for one generation and collected adults from the second generation for DNA extraction.

For each species, we extracted from one individual using an adjusted version of the 10X Genomics sample preparation protocol “DNA Extraction from Single Insects” (https://assets.ctfassets.net/an68im79xiti/3oGwQ5kl6UyCocGgmoWQie/768ae48be4f99b1f984e21e409e801fd/CG000145_SamplePrepDemonstratedProtocol_-DNAExtractionSingleInsects.pdf). DNA concentrations were measured with a Qubit 3.0 Fluorometer using the dsDNA HS Assay Kit (Thermo Fisher Scientific) and DNA integrity was assessed on an agarose gel stained with 2% GelRed. 10X Genomics Chromium linkedread sequencing libraries were prepared and subsequently sequenced to yield paired-end 2×150 bp reads, on a HiSeq X platform at SciLifeLab (Stockholm, Sweden).

### Genome assembly and scaffolding

Raw 10X genomics reads were checked for sequencing quality using FastQC v0.11.5 (Andrews 2010), and *de novo* assembled using the Supernova 2.0 (Weisenfeld *et al.* 2017) assembler. We then polished the draft assembly using purge_dups (Guan *et al.* 2020) to remove haplotigs and heterozygous overlaps based on sequence similarity and read depth. Subsequently, assemblies were scaffolded using arcs (Yeo *et al.* 2018) and links (Warren *et al.* 2015). To remove sequence contamination from the assembly, we ran Kraken v2.0 (Wood and Salzberg 2014) against bacterial, archaeal, and viral domains, along with the human genome. We assessed the completeness of our polished genome assemblies assessed by Benchmarking Universal Single-Copy Orthologs (BUSCO) from OrthoDB v1.22 (Simão *et al.* 2015) using Endopterygota as the taxonomic database.

### Gene annotation

We first assessed the repeat content of our genome assemblies and created a specific repeat library using RepeatModeler 1.0.11 (Smit *et al.* 2010b) for each genome assembly. Based on the repeat library, identification of repeat sequences in the genome was performed using RepeatMasker 3.0.9 (Smit *et al.* 2010a) and RepeatRunner (Yandell 2006) with default settings. RepeatRunner is a program that integrates RepeatMasker with BLASTX, allowing the analysis of highly divergent repeats and identifications of divergent protein-coding portions of retro-elements and retroviruses.

Gene annotation was performed using the MAKER package v3.01.02 (Holt and Yandell 2011). First, for each genome, we generated one initial evidence-based annotation using both protein and transcriptome data sources. Protein databases came from the Uniprot Swiss-Prot database (downloaded on 2019-11; 561,356 proteins) (Engler *et al.* 2020), as well a subset of manually selected proteins (uniport request: taxonomy: “Coleoptera [7041]”, existence: “Inferred from homology [3]”, 161,853 proteins). In addition to protein resources, transcriptome data containing 57,255 transcripts from *G. pusilla* were used as evidence for all three genomes (Yang *et al.* 2020). Next, we used the candidate genes from the initial annotation to train three different *ab-initio* gene predictors: GeneMark v4.48 (Besemer *et al.* 2001), Augustus v3.3.3 (Stanke *et al.* 2006) and Snap v2013_11_29 (Korf 2004). Finally, an *ab-initio* evidence-driven gene build was generated based on the initial evidence-based annotation and the *ab-initio* predictions. Additionally, we used EVidenceModeler v 1.1.1 (Haas *et al.* 2008), which allows the construction of gene models based on the best possible set of exons produced by the *ab-initio* tools, and chooses those most consistent with the evidence. Functional inference for genes and transcripts was performed using the translated CDS features of each coding transcript.

Each predicted protein sequence was run against InterProscan (Jones *et al.* 2014) in order to retrieve functional information from 20 different sources. In addition, Blastp was performed against the complete Swissprot/Uniprot database (downloaded 2019-11) with a maximum e-value cut-off of 1*e*-6 to assign putative functions to predicted proteins. tRNA have been predicted through tRNAscan v 1.3.1 (Lowe and Eddy 1997).

### Ortholog cluster analysis

Identifying shared orthologous clusters allows the comparison of function and evolution of proteins across closely related species. An ortholog cluster analysis was performed by comparing the three complete *Galerucella* protein sets with each other via OrthoVenn2 with default settings of E = 1*e*-5 and an inflation value of 1.5.

### Data availability

Raw read data, the final assemblies and annotations are available at the EMBL-ENA database under BioProject PRJEB44256. Supplemental material available at figshare: https://figshare.com/account/home#/projects/111680

## Results and discussion

### Genome assemblies

Sequencing of the 10X genomics libraries yielded a total of 683.34 million read pairs, resulting in a sequencing depth above 110X for each species. Due to the low molecular weight of the input DNA, the initial *de novo* assembly from Supernova was highly fragmented, with N50 values of 49.884kb, 19.764kb and 24.604kb for *G. calmariensis*, *G. pusilla* and *G. tenella*. Redundancy removal by purge_dups and arcs+links scaffolding dramatically improved the quality and contiguity of assemblies (see supplementary materials Table 1 for the comparisons between assemblies). The decontamination process removed two contigs from *G. calmariensis*, one contig from *G. pusilla* and two contigs from *G. tenella* which matched the human database with a kmer length >100bp. Final assemblies for *G. calmariensis* had a size = 588Mb, contained 39,255 scaffolds with a N50 = 79.674kb, final assemblies for *G. pusilla* had a size = 513Mb, 40,929 scaffolds with a N50 = 45,442kb whereas final assemblies for *G. tenella* has a size = 460Mb, 29,202 scaffolds with a N50 = 31,588kb (Table 1). Although the final assembly was still fragmented, the completeness of genome measured by BUSCO was satisfactory. Using 2124 BUSCO groups with endopterygota_odb10 database, we found 91.3% complete orthologs and only 4.0% missing orthologs in *G. calmariensis*, 85.3% complete orthologs and 6.5% missing orthologs for *G. pusilla* and 95.4% complete orthologs and 3.3% missing orthologs for *G. tenella*. The GC content of the three genomes ranged from 33.6% to 33.8%, which is slightly higher than the GC content of the ragweed leaf beetle genome assembly (Bouchemousse *et al.* 2020).

**Table 1.**
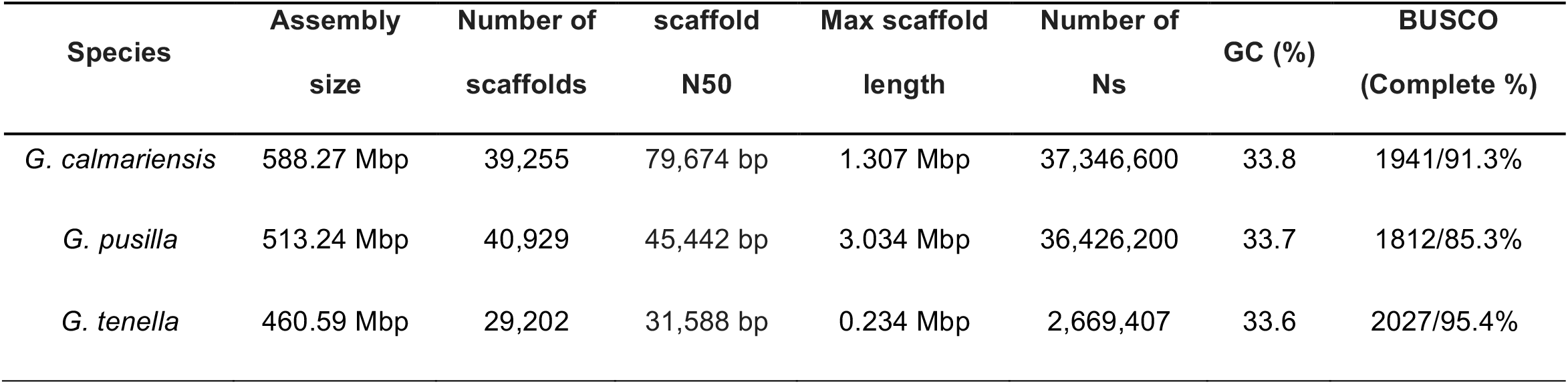
Summary of *G. calmariensis*, *G. pusilla* and *G. tenella* reference genomes. BUSCO score is based on the Endopterygota_db10 dataset.

The sizes of the genome assemblies of our three *Galerucella* species varied (460Mb to 588Mb) but is slightly smaller than the size of the Colorado potato beetle (642 Mb) (Herndon *et al.* 2020) and the Ragweed leaf beetle (774 Mb) (Bouchemousse *et al.* 2020). Coleoptera is amongst the most diverse insect orders in terms of genome size, with an average genome size of 760MB and ranges from 160 to 5,020 Mb (Gregory 2021). Within-genus variation in genome size is relatively small in these three assemblies compared with other Coleopteran species, possibly because of their close phylogenetic relationships and similarities in life cycle, food sources and wasp enemies.

#### Gene annotation

RepeatMasker masked 48.55%, 46.65% and 40.84% of the *G. calmariensis*, *G. pusilla* and *G. tenella* genomes as repetitive elements. In addition, RepeatRunner further masked approximately 1% of each genome as repeats using MAKER TE as library (Supplementary table 2).

The *ab initio* evidence-driven annotation using the MAKER pipeline revealed 32,294, 30,302 and 33,794 potential protein coding genes, accounting for 16.2%, 17.7% and 19.1% of the whole genome of *G. calmariensis*, *G. pusilla* and *G. tenella* respectively (Supplementary 3). For each species, 84% to 86% of protein coding genes were assigned with a putative function, and 39% to 45% had a GO annotation (supplementary table 4, functional annotations using InterProscan from 20 different sources). Blast against the Uniprot/Swissprot database predicted 15,046, 14,404 and 17,958 hits with unique gene names for *G. calmariensis*, *G. pusilla* and *G. tenella*, respectively.

#### Ortholog cluster analysis

The three protein sets of *Galerucella* species were compared to identify shared orthologous clusters using OrthoVenn2 (Figure 2). The complete protein sets contain 40,031 sequences from *G. calmariensis*, 37,514 sequences from *G. pusilla* and 44,200 sequences from *G. tenella*, corresponding to 20,665, 19,730 and 19,106 ortholog clusters respectively.

**Figure 2.**
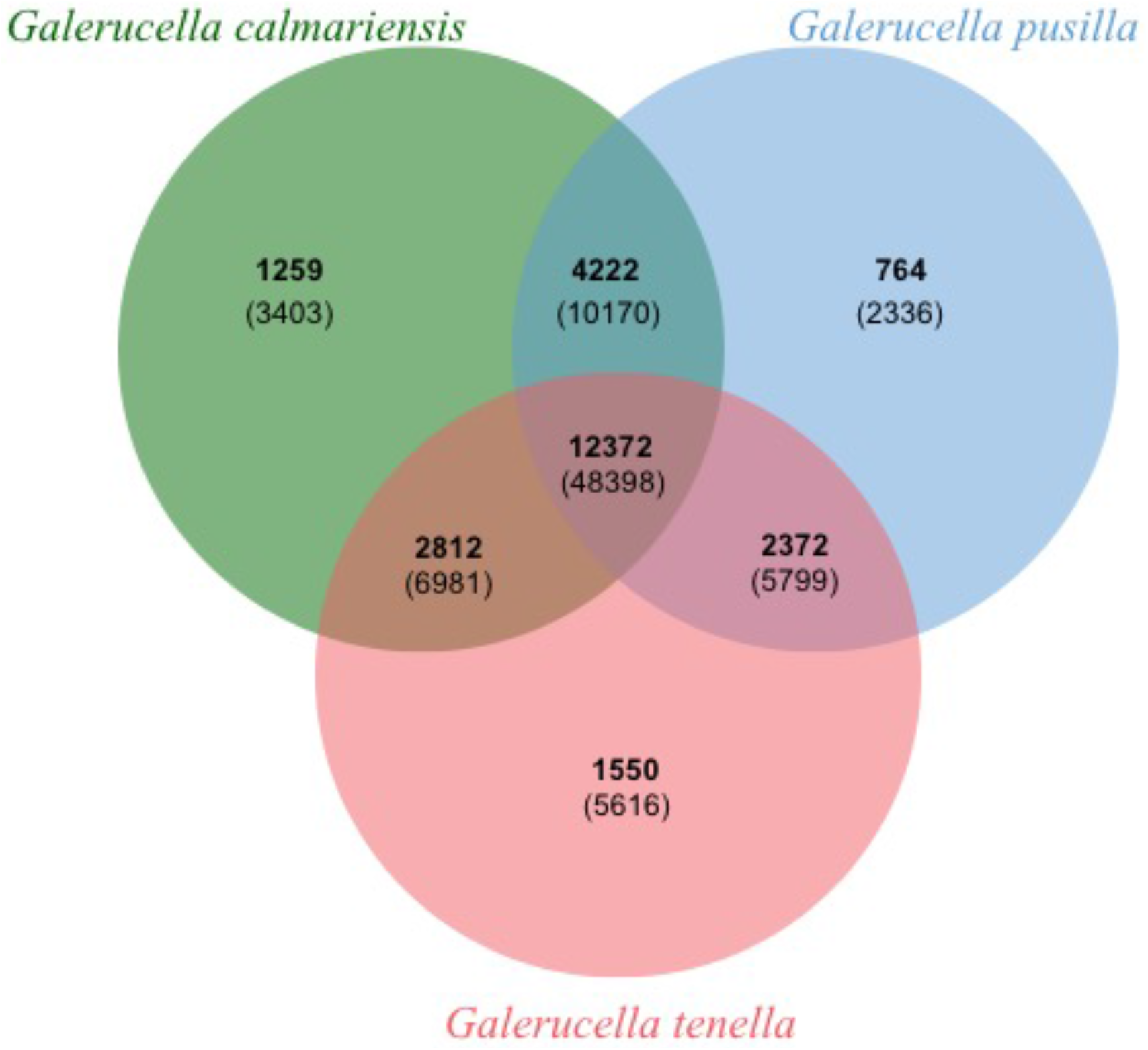
A Venn diagram of the orthologous gene clusters among the three *Galerucella* species: *G. calmariensis*, *G. pusilla* and *G. tenella*. The numbers of shared Ortholog clusters between species is indicated in the overlapping areas of the circles while the numbers of proteins corresponding to each cluster are underneath in parentheses

Most annotated genes (12,372 orthogroups/48,398 proteins) were shared between the three species. Shared clusters between *G. calmariensis* and *G. pusilla* (16,594) account for 80.3% and 84.1% of ortholog clusters in *G. calmariensis* and *G. pusilla* respectively whereas the shared regions of either *G. calmariensis* and *G. pusilla* with *G. tenella* account for less than 75% of their clusters. Ortholog clusters unique to a single species account for 6.09%, 3.87% and 8.11% of the entire cluster set for *G. calmariensis*, *G. pusilla* and *G. tenella*, which indicates divergent regions between species (Ferguson *et al.* 2020). The inflated numbers of singleton clusters in *G. tenella* may be due to the high duplication levels in the genome, as BUSCO detected 33.1% complete duplicated BUSCOs in *G. tenella*. Whether this is due to gene duplication or assembly error should be further investigated. The duplication and fragmentation level detected by BUSCO are similar between *G. calmariensis* (1.8% duplicated and 5.1% fragmented) and *G. pusilla* (1.2% duplicated and 8.2% fragmented), however *G. calmariensis* harbours a higher level of singleton clusters than *G. pusilla*.

## Conclusion

*Galerucella* species plays an important role as biological control agents as well as for ecological and evolutionary research. Here we produced draft genome assemblies for three leaf beetles in the *Galerucella* genus, which are the first three genomes from the Galerucinae subfamily branch of the leaf beetle family. These draft genomes will be valuable for investigating a deeper resolution in phylogenetic comparisons of Coleoptera. In addition, the genomes and annotations will provide a valuable resource for the monitoring of biocontrol methods. The genome sequencing of the three closely related beetles sharing a common wasp enemy also provide possibilities of understandings of food web and host-parasitoid interactions. In particular, comparing genomes of species with divergent immune resistance against parasitoid wasps helps detecting essential genetic regions underlying host immunity and other potential traits participating in the arms race between host and parasitoids.

## Acknowledgements

The authors would like to acknowledge support from Science for Life Laboratory, the National Genomics Infrastructure, NGI, and Uppmax for providing assistance in massive parallel sequencing and computational infrastructure. Computational analyses were enabled by resources provided by the Swedish National Infrastructure for Computing (SNIC) through Uppsala Multidisciplinary Center for Advanced Computational Science (UPPMAX) under project snic2017-7-233. SNIC is partially funded by the Swedish Research Council through grant agreement no. 2018-05973”.

## Footnotes

Supplemental material available at figshare: (private link) https://figshare.com/s/49c775f1be0691fdcbc5

